# Decrease in dose per fraction impairs the FLASH sparing effect in murine intestine model

**DOI:** 10.1101/2025.08.27.672569

**Authors:** A. Sesink, R. Geyer, P. Devanand, T. T. Böhlen, L. Soutter, R. Moeckli, C. Bailat, F. G. Herrera, V. Grilj

## Abstract

**Purpose:** FLASH radiotherapy (FLASH) can ease radiation-induced normal tissue toxicities; however, its benefit in clinically relevant fractionation protocols remains insufficiently explored. This study investigated the FLASH sparing effect under two fractionated regimens using a murine model of acute gastrointestinal toxicity.

**Methods and Materials:** Tumor-free C57BL/6 mice received abdominal irradiation with either conventional radiotherapy (CONV) or FLASH using a 9 MeV electron beam. Three dose delivery protocols were assessed: single-fraction delivery, two equal fractions over two consecutive days, and ten equal daily fractions over two weeks. Dose escalation was performed within each protocol, while overall survival was used to monitor normal tissue sparing. The FLASH dose modifying factor (FDMF) was derived from normal tissue toxicity probability (NTCP) curves to quantify the relative protective effect of FLASH.

**Results:** The single-fraction irradiation demonstrated a significant FLASH sparing effect, with an FDMF of 1.14. In contrast, this protective effect was diminished in the fractionated protocols, with the two-fraction regimen yielding an FDMF of 1.03, while the ten-fraction regimen showed no measurable sparing (FDMF = 1.00).

**Conclusions:** In an acute responding model of radiation-induced abdominal toxicity, the FLASH sparing effect was substantially reduced with a two-fraction regimen and completely absent with a ten-fraction regimen. These findings suggest that the benefit of FLASH may be limited at lower doses per fraction and highlight the need for further studies in other clinically relevant models to better define the boundaries of its therapeutic applicability.

**Highlights:**

- FLASH-RT spares mice intestine from acute toxicity for single-dose delivery.
- FLASH effect declines in mice intestine with more fractions and less dose/fraction.
- FLASH sparing in mice intestine is lost with ten equal fractions over two weeks.

## INTRODUCTION

Radiation therapy plays an essential role in cancer treatment, with more than 50% of patients worldwide receiving it at some stage of their care (1). Over the past years, major technological advancements, have significantly improved precision and targeting of tumors. However, dose fractionation continues to be a critical element for maximizing the tumor control, while minimizing the adverse effects on healthy tissues (2).

Administering radiation in smaller doses over several weeks or months enables reoxygenation of hypoxic cancer cells and redistribution of cancer cells into more radiosensitive phases of the cell cycle, increasing their susceptibility to radiation. Concurrently, this approach promotes repair of sublethal damage in healthy tissues, thereby reducing radiation-induced side effects (3, 4). Although fractionation offers several advantages, the total dose that can be delivered is still constrained by the tolerance of surrounding healthy tissues, restricting tumor control and overall effectiveness of the treatment (5, 6).

FLASH radiotherapy (FLASH), characterized by ultra-high dose rates (UHDR > 100 Gy/s) (7-13), has emerged as a promising approach in radiation oncology for mitigating the dose limitations of conventional radiotherapy (CONV). In preclinical studies this innovative radiation delivery, administered within milliseconds, demonstrated to significantly reduce normal tissue toxicity while maintaining comparable anti-tumor efficacy to CONV delivered over minutes (8, 11, 12, 14-19). However, the FLASH sparing effect has been predominantly demonstrated in animal models for single-fraction irradiations of large doses using proton or electron beams (8, 11, 15, 16, 18, 20-25). The observed FLASH dose modifying factors (FDMF) ranged from 1.1 in case of the intestine models up to 1.45-1.50 for skin (25, 26). For FLASH to be successfully integrated into clinical practice, it is essential to demonstrate that the requirements for eliciting the FLASH sparing effect are not in collision with standard dose delivery protocols including dose fractionation.

Recent studies suggest that the FLASH sparing effect in normal brain tissue can be achieved with hypofractionated radiotherapy of 2-3 fractions (27, 28) and following a standard-of-care protocol comprising 10 daily fractions of 3 Gy (29). However, these findings have not been validated in other tissues and for different endpoints, leaving the broader reproducibility of the FLASH sparing effect under fractionated delivery still uncertain. Moreover, the magnitude of the FLASH sparing effect, quantified as the ratio of isoeffective doses between FLASH and CONV, has not yet been studied in the context of fractionated FLASH treatments.

In this study, we aimed to investigate the reproducibility of the FLASH sparing effect with two hypofractionated protocols (two fractions over two consecutive days and ten fractions over two consecutive weeks) using a model of radiation-induced acute gastrointestinal toxicity.

## MATERIALS AND METHODS

### Irradiation setup

All irradiations were carried out using the FLASH Mobetron LINAC (IntraOp, Sunnyvale, CA, USA), with the machine’s commissioning previously described by Moeckli et al. (2021) (30). Animals were placed in a supine position on a water-equivalent RW3 slab phantom (PTW Freiburg, Germany) and stabilized laterally by two 15 mm-thick RW3 blocks. The blocks were spaced approximately 2 cm apart, with the distance adjusted individually for each mouse to ensure the abdominal surface was level with the top surface of the blocks as represented in (**Figure 1a** and **1b**). Whole-abdomen irradiations were administered with 9 MeV electron beam at either CONV dose rates (0.5 Gy/s) or ultra-high dose rates characteristic for FLASH (> 200 Gy/s). A circular 3 cm-diameter collimator, attached to a 20 cm long cylindrical applicator, was positioned in contact with the two RW3 blocks to deliver the radiation. Animals were manually aligned with the collimator, ensuring that the resulting radiation field covered the entire abdominal region.

**Figure 1.**
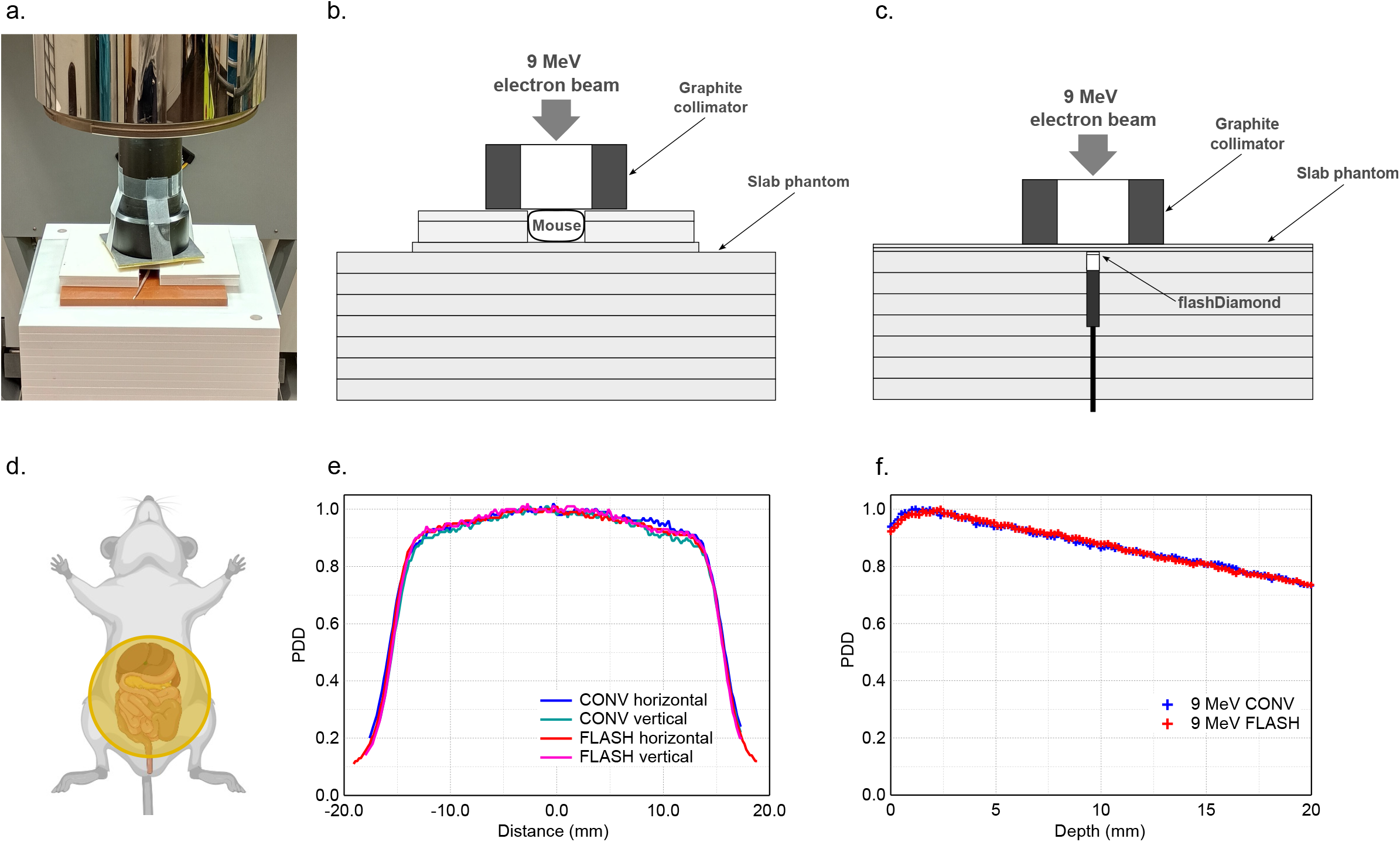
Irradiation setup for whole abdomen irradiations. (a) Photograph of the animal being irradiated by Mobetron accelerator using a circular carbon collimator of 3 cm diameter supported by a 25 cm long applicator tube. Schematic illustrations of the setup for (b) mouse irradiations, and (c) reference dosimetry using flashDiamond. (d) Illustration of animal positioning with the exposed abdominal cavity region in correspondence with the radiation field. (e) Horizontal (CONV, blue; FLASH, red) and vertical (CONV, green; FLASH, pink) dose profiles measured with EBT3 films at the surface of a solid water phantom. (f) Percent depth dose (PDD) profiles measured following CONV (blue) and FLASH (red) in water.

### Fractionation regimens

Three irradiation protocols were employed based on the timing of dose delivery: a single-fraction protocol, in which the entire dose was administered in one uninterrupted session; a two-fraction protocol, with two equal doses delivered over two consecutive days (24-hour interval); and a ten-fraction protocol, involving ten equal doses delivered Monday through Friday across two weeks, with a break over the weekend. The dose ranges were as follows: 14–17 Gy for single-fraction CONV and 14–20 Gy for single-fraction FLASH treatments; 8–13 Gy for the two-fraction protocol; and 5–7 Gy for the ten-fraction protocol. The specific CONV and FLASH beam parameters are listed in **Supplementary tables S1–S4**.

### Dosimetry

Radiation doses were prescribed at a depth of 5 mm in the water-equivalent RW3 slab phantom and measured before and after animal irradiations with the flashDiamond detector (T60025, PTW Freiburg, Germany) (**Figure 1c**). Signal acquisition from the detector was performed using a UNIDOS E electrometer (PTW, Freiburg, Germany). To enhance measurement redundancy, a GafChromic EBT3 film (Ashland, Bridgewater, NJ, USA) was simultaneously irradiated at a depth of 4 mm alongside the flashDiamond detector. The uncertainty related to flashDiamand detector was ± 0.5% (31). Film analysis was performed as described by Jaccard M. et al. (2017) (32). Globally, the dose uncertainty on each mouse irradiation was estimated to be below 3%. Results of dose verification measurements by the flashDiamond detector for single-, two- and ten-fraction experiments are presented in **Supplementary tables S5-S7**. CONV and FLASH beams produced using the 3 cm collimator were initially characterized during machine commissioning (30). However, we re-assessed their lateral and depth dose profiles using the EBT3 films to ensure a reliable comparison of their respective biological effects (**Figure 1e–f**).

### Animal handling

All animal experiments within this research were approved and performed in accordance with the guidelines established by the Animal Ethics Committee of Canton Vaud, Switzerland (Approval nr. VD3962a). Mice used within this research were acclimated for 1 week prior to experimentation. Mice were maintained and handled in pathogen-free conditions in cages with a maximum of five mice per cage, under a controlled 12h light/dark cycle, relative humidity of 55%, controlled temperature of 21 °C, with food and sterile water provided daily, all in accordance with the guidelines provided by the institution. All experiments were conducted on female C57BL/C mice (Charles River; France) at 8-10 weeks of age. Prior to irradiation, mice were placed under ketamine anaesthesia (75 µg/g body weight), supplemented with Medetomidine (0.5 µg/g body weight) using intraperitoneal (IP) injection and followed by Atipamezole (1 µg/g body weight) IP injection post-irradiation.

### Survival analysis

Normal tissue toxicity probability (NTCP) is expressed as the number of animals euthanized because of gastrointestinal toxicity, in compliance with the approved animal license. Survival times were calculated from the time of irradiation until euthanasia. Symptoms of acute toxicity (including weight loss, hunching, diarrhea, activity, behavioral changes), were monitored for up to 15 days following the initial irradiation for a single and two-fraction regimens, and up to 25 days for the ten-fraction regimen. These symptoms were documented and scored to determine whether early euthanasia criteria were met (i.e. permanent body weight loss > 15%; or severely decreased activity; or combination of reduced activity, bodyweight loss > 10% and diarrhea). Animals were euthanized via CO2 asphyxiation followed by secondary cervical dislocation at the 15- or 25-day endpoint unless euthanasia was required earlier.

### FLASH dose modifying factor

For each fractionation regimen, the FLASH dose-modifying factor (FDMF) was determined by calculating the ratio of LD_50_ values (the doses corresponding to a 50% NTCP) between FLASH and CONV treatments, based on probit fits of NTCP versus dose data.

### Statistical analysis

Reported doses for each group represent the mean dose calculated across all animals within that group. NTCP curves were generated using probit regression fits based on individual mouse data, with proportions displayed along the fit and error bars representing Jeffreys 68% (1-sigma) credible intervals for binomial proportions. Survival analysis was performed using the Kaplan-Meier method. The log-rank (Mantel–Cox) test was used to compare FLASH and CONV survival curves in all groups where equal dose delivery resulted in differing overall survival rates. The statistical analyses were performed using GraphPad Prism (v 7.03). Statistical significance was set at *P < 0.05, **P < 0.01, ***P < 0.001 and, ****P < 0.0001.

## RESULTS

### FLASH mitigates acute intestine toxicity following single fraction delivery

To confirm that our FLASH beam reduces acute intestinal toxicity in a single-fraction regimen, we conducted whole-abdomen irradiations on female C57BL/6 mice with escalating doses of FLASH and CONV (**Figure 2a**). The resulting NTCP values are plotted as a function of dose in **Figure 2b**. The LD_50_-derived FDMF was 1.14, suggesting that a 14% higher single-fraction dose of FLASH is needed to elicit the same level of toxicity as CONV. Abdominal irradiation with 14 Gy CONV, 14 Gy FLASH, and 16 Gy FLASH resulted in 15-day overall survival rates above 95%. At doses above 16 Gy, all animals irradiated by CONV required euthanasia, whereas FLASH at 17 and 18 Gy yielded 15-day survival rates of 50% and 20%, respectively (**Figure 2c-d**). Comparative survival analyses of CONV and FLASH at 16 and 17 Gy yielded p-values of <0.0001 and 0.0072, respectively (**Supplementary Figure 1**). Median survival times for all single-fraction groups are listed in **Supplementary Table 8**.

**Figure 2.**
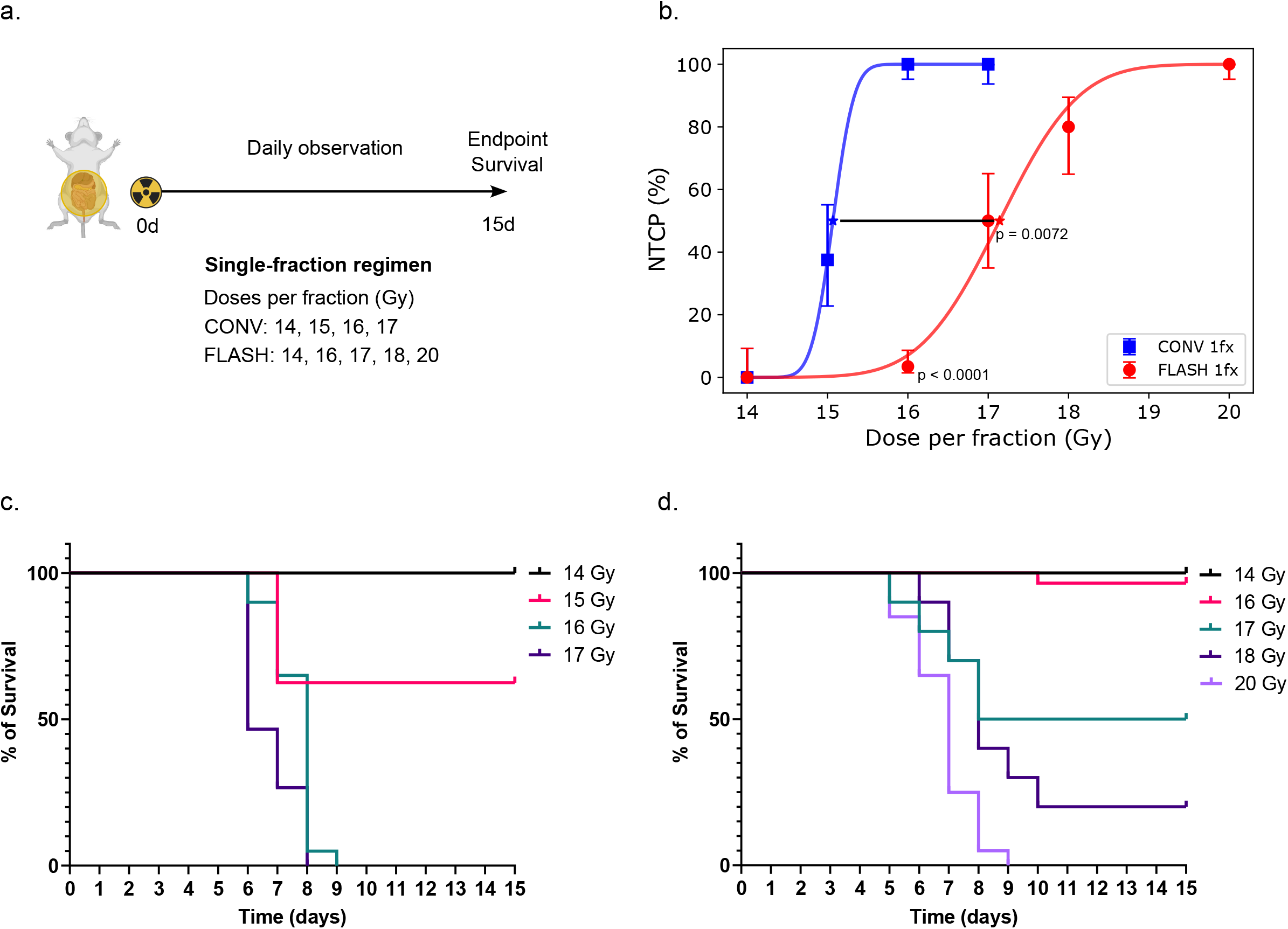
FLASH mitigates acute intestine toxicity following single fraction delivery. Experimental scheme (a) and obtained NTCP values (b) for single-fraction delivery of escalating doses of FLASH and CONV. CONV doses included 14 (n = 10), 15 (n = 8), 16 (n = 20), and 17 Gy (n = 15), while FLASH doses were 14 (n = 10), 16 (n = 29), 17 (n = 10), 18 (n = 10) and 20 Gy (n = 20). Stars indicate LD_50_ values obtained from probit fits (CONV, blue; FLASH, red) of NTCP data. Error bars represent estimated uncertainty using Jeffreys 1-sigma credible intervals. The indicated p-values were calculated by comparing CONV and FLASH survival curves at matched dose levels using the log-rank test. (c)-(d) Kaplan-Meier survival curves showing 15-day outcomes following single-fraction whole-abdomen irradiation with CONV and FLASH. The survival rates observed for CONV were 100% at 14 Gy, 62.5% at 15 Gy, and 0% at 16 Gy and 17 Gy. The survival rates observed for FLASH were 100% at 14 Gy, 95.6% at 16 Gy, 50% at 17 Gy, 20% at 18 Gy and 0% at 20 Gy.

### Attenuation of the FLASH sparing effect under two-fraction delivery

After confirming the FLASH sparing effect with single-fraction irradiation, we examined its reproducibility under different hypofractionated protocols. The two-fraction regimen consisted of delivering two equal doses separated by a 24-hour interval. (**Figure 3a**). Escalating doses per fraction (8 – 13 Gy) were administered using both CONV and FLASH. The resulting NTCP are presented as a function of dose per fraction in **Figure 3b**. Although there was a slight trend suggesting reduced abdominal toxicity with FLASH, the difference was not statistically significant. Based on LD_50_ values derived from probit fits of CONV and FLASH datasets, the calculated FDMF was 1.03.

**Figure 3.**
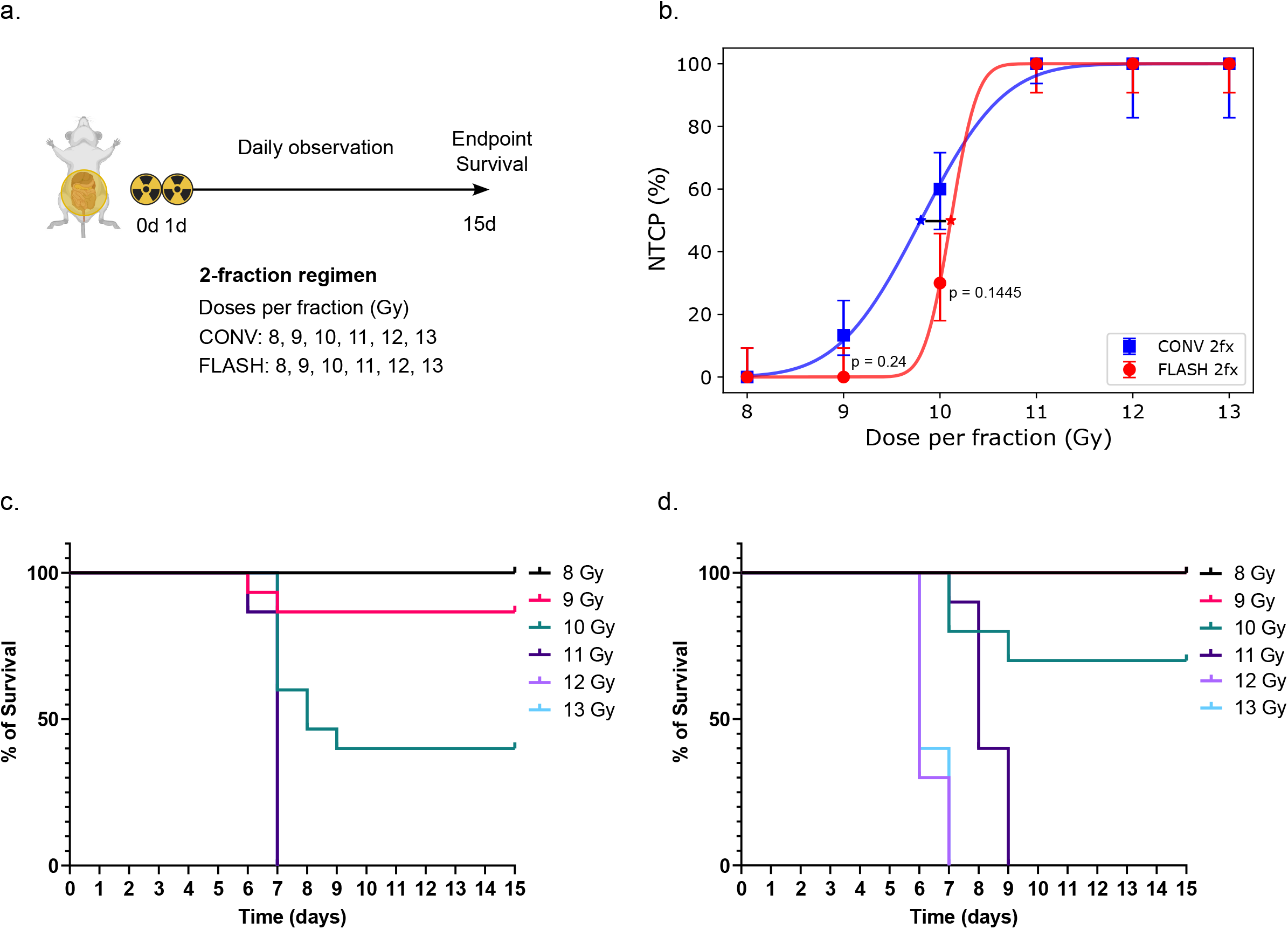
Attenuation of the FLASH sparing effect under two-fraction delivery. Experimental scheme (a) and obtained NTCP values (b) for two-fraction delivery of escalating doses of CONV and FLASH. For CONV, doses per fraction of 8 (n = 10), 9 (n = 15), 10 (n = 15), 11 (n = 15), 12 (n = 5), and 13 (n = 5) Gy were used, while for FLASH, all dose levels from 8 to 13 Gy were tested with n = 10 per group. Stars indicate LD_50_ values obtained from probit fits (CONV, blue; FLASH, red) of NTCP data. Error bars represent estimated uncertainty using Jeffreys 1-sigma credible intervals. The indicated p-values were calculated by comparing CONV and FLASH survival curves at matched dose levels using the log-rank test. (c)-(d) Kaplan-Meier survival curves showing 15-day outcomes following two-fraction irradiations of whole-abdomen with CONV and FLASH. The survival rates observed for CONV were 100% at 8 Gy/fraction, 86.7% at 9 Gy/fraction, 40% at 10 Gy/fraction, 0% at 11 Gy/fraction, 12 and 13 Gy/fraction. The survival rates observed for FLASH were 100% at 8 and 9 Gy/fraction, 70% at 10 Gy/fraction, and 0% at 11, 12 and 13 Gy/fraction.

Administration of two fractions of 8 Gy CONV, 8 Gy FLASH, and 9 Gy FLASH resulted in a 100% overall survival rate at 15 days after the first dose (**Figure 3c-d**). Two fractions of 9 Gy CONV yielded a slightly lower survival rate of 87%; however, this difference compared to the 2 × 9 Gy FLASH group was not statistically significant (p = 0.24). Among animals receiving two fractions of 10 Gy, survival at day 15 was 40% in the CONV group and 70% in the FLASH group, though this difference also did not reach statistical significance (p = 0.1445). Groups exposed to two fractions of 11 Gy or higher required euthanasia regardless of the beam modality. Survival outcomes for two-fraction CONV and FLASH treatments at 9 and 10 Gy per fraction are compared in **Supplementary Figure 2**. The median survival times for all two-fraction groups are provided in **Supplementary Table 9**.

### Loss of FLASH sparing effect in ten-fraction delivery of sub-8 Gy doses

The ten-fraction regimen consisted of administering ten equal fractions of CONV and FLASH over a two-week period, with dose escalation from 5 to 7 Gy per fraction for both modalities (**Figure 4a**). The resulting NTCP values showed no difference in acute abdominal toxicity between CONV and FLASH treatments, indicating a complete absence of the FLASH sparing effect under this fractionation regimen (**Figure 4b**). This is further evidenced by an FDMF at LD_50_ of 1.00. Ten fractions of 5 Gy led to a 100% overall survival rate at 25 days following the first dose for both CONV and FLASH treatments, while doses of 6.5 Gy or 7 Gy per fraction led to incomplete treatment administration, with all animals requiring euthanasia during the second week of irradiation (**Figure 4c-d**). At 6 Gy per fraction, severe toxicity symptoms began to appear during the second week only in the FLASH group, but the difference in survival curves was not statistically significant (p = 0.897). Both regimens resulted in 0% survival within five days following the final fraction. Comparative survival analyses of these two groups is shown in **Supplementary Figure 3**. At 5.5 Gy per fraction, the 25-day survival rate was 80% in the CONV group and 75% in the FLASH group (p = 0.6214). The median survival times for all 10-fraction groups are given in **Supplementary Table 10**.

**Figure 4.**
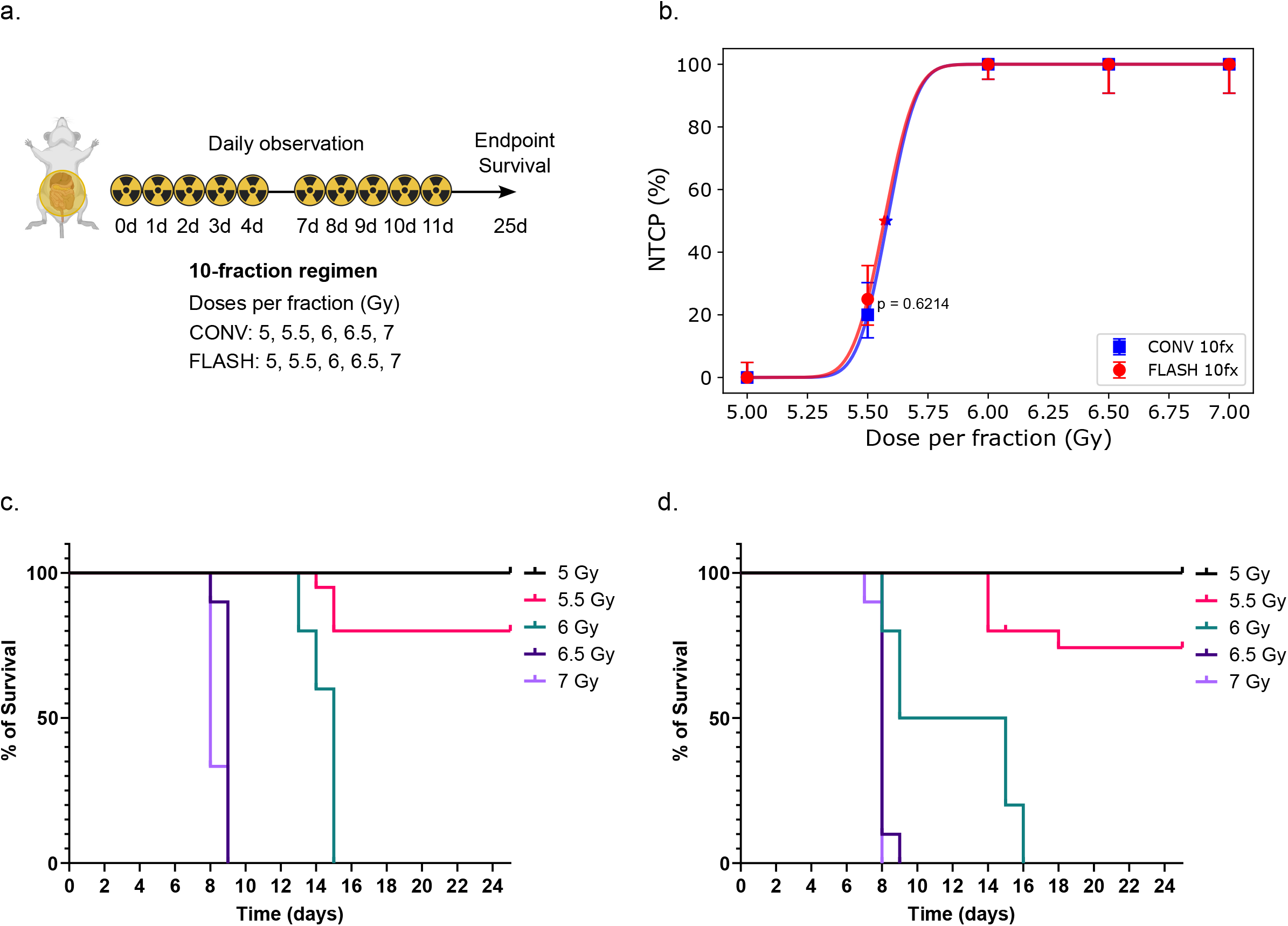
Loss of FLASH sparing effect in ten-fraction delivery of sub-8 Gy doses. Experimental scheme (a) and obtained NTCP values (b) for ten-fraction delivery of escalating doses of CONV and FLASH. Doses per fraction of 5 (n = 20), 5.5 (n = 20), 6 (n = 20), 6.5 (n = 10) and 7 (n = 10) Gy were used for both CONV and FLASH. Stars indicate LD_50_ values obtained from probit fits (CONV, blue; FLASH, red) of NTCP data. Error bars represent estimated uncertainty using Jeffreys 1-sigma credible intervals. The indicated p-values were calculated by comparing CONV and FLASH survival curves at matched dose levels using the log-rank test (c)-(d) Kaplan-Meier survival curves showing 25-day outcomes following ten-fraction whole-abdomen irradiation with CONV and FLASH. The survival rates observed for CONV were 100% at 5 Gy/fraction, 80% at 5.5 Gy/fraction, 100% at 6 Gy/fraction, and 0% at 6.5 and 7 Gy/fraction. The survival rates observed for FLASH were 100% at 5 Gy/fraction, 75% at 5.5 Gy/fraction, 100% at 6 Gy/fraction, and 0% at 6.5 and 7 Gy/fraction.

## DISCUSSION

The potential to maintain the FLASH sparing effect with dose fractionation remains largely unexplored. Fractionated FLASH treatment has been previously studied only in the mouse brain; however, the influence of time fractionation schedules on the FLASH sparing effect in early-responding organs has not been investigated. The primary objective of our study was to evaluate whether the FLASH sparing effect is preserved in an abdominal model of acute radiation induced toxicity under two different hypofractionated protocols. To achieve this, we delivered escalating whole-abdomen doses in a single-, two- and ten-fraction regimens, with overall survival post-irradiation as the primary endpoint.

We demonstrated that delivering a single fraction of FLASH yielded an FDMF of 1.14, significantly enhancing survival outcomes compared to CONV and surpassing previously reported FDMF values in this model (26). When dose was delivered in two fractions, a non-significant trend suggesting some degree of FLASH sparing persisted; however, the FDMF was reduced to 1.03. With ten fractions, there was no evidence of the FLASH sparing effect, as indicated by an FDMF of 1.00. The results obtained are in contrast with earlier studies reporting a preserved FLASH sparing effect after two, three, and ten fractions (10, 27-29, 33), though it is important to note that such studies evaluated late toxicity at a single dose level. In this case, calculation of the FDMF is not feasible, leaving it unclear whether the extent of FLASH-induced normal tissue sparing is reduced under fractionated delivery compared to single-dose treatment in the same model. Moreover, unlike our study, the findings for the 10 times 3 Gy fractionation regimen were based on a highly specific ex vivo endpoint that lacks functional relevance (29).

An explanation for the reduced sparing capacity seen with fractionated FLASH treatments might be in the potential dose dependence of the FLASH sparing effect. This relationship, previously suggested by Böhlen et al. (24), likely varies across tissue types and endpoints, implying that minimum dose per fraction may be necessary to preserve normal tissue protection under fractionated FLASH regimens. In our murine abdominal model, the 10-fraction data indicate that, if such a dose requirement exists, it likely exceeds 6 Gy. Further research into how the FLASH sparing effect varies with dose across tissues and endpoints is essential for accurately assessing the clinical benefit of FLASH treatment. If achieving the FLASH sparing effect requires high doses per fraction and extremely hypofractionated regimens, the loss of the protective benefits provided by conventional fractionation could offset any therapeutic gains (34). Exploring the impact of inter-fraction interval on normal tissue response to FLASH is another important area for future investigation. For example, employing 48-hour intervals, as proposed in ultra-hypofractionated RT (35) or stereotactic body radiation therapy (SBRT) (36) for prostate cancer, could produce different outcomes compared to the 24-hour intervals used in this study. Whether acute radiation toxicity is reduced with FLASH under such conditions remains unclear. All these considerations are crucial for informing the safe and effective clinical translation of FLASH-RT.

Additionally, the gastrointestinal toxicity model used showed a comparatively lower FDMF than brain and skin models (10, 16, 18, 24). It is possible that fractionation may have a smaller impact on the magnitude of the FLASH sparing effect in these other models with higher FDMF values or in late toxicities compared to early ones. Nevertheless, our findings support the suitability of this model as a rapid *in vivo* screening tool for detecting the presence or absence of the FLASH sparing effect.

## CONCLUSION

This research provides evidence that in a model of radiation-induced acute abdominal toxicity, the FLASH sparing effect observed with single-fraction delivery, is adversely affected by dose fractionation. The FDMF decreased from 1.14 with a single fraction to 1.03 with two fractions, and further to 1.00 with a ten-fraction regimen, highlighting the potential difficulty of maintaining the conditions necessary for the FLASH sparing effect in early-responding tissues under fractionated protocols. Further research into the potential dose dependence of the FLASH sparing effect is needed to better define the limitations of FLASH and accurately evaluate its clinical applicability.

### Declaration of Generative AI and AI-assisted technologies in the writing process

*During the preparation of this work the author(s) used ChatGPT in order to improve the readability and language. After using this tool/service, the author(s) reviewed and edited the content as needed and take(s) full responsibility for the content of the publication*

## Acknowledgements

This work was supported by ISREC Foundation and Swiss National Science Foundation (project number 215766). We are grateful to the staff at Agora and Epalinges animal facilities for their expertise in animal handling and maintenance throughout this study.

## Bibliography

1. Lievens Y, Borras JM, Grau C. Provision and use of radiotherapy in Europe. Mol Oncol. 2020;14(7):1461–9.

2. Meattini I, Becherini C, Boersma L, Kaidar-Person O, Marta GN, Montero A, et al. European Society for Radiotherapy and Oncology Advisory Committee in Radiation Oncology Practice consensus recommendations on patient selection and dose and fractionation for external beam radiotherapy in early breast cancer. Lancet Oncol. 2022;23(1):e21–e31.

3. Hall E, Giaccia AJ. Radiobiology for the Radiologist. 8 ed: Wolters Kluwer; 2018.

4. van Aken ESM, Devnani B, Castelo-Branco L, Ruysscher D, Martins-Branco D, Marijnen CAM, et al. ESMO-ESTRO framework for assessing the interactions and safety of combining radiotherapy with targeted cancer therapies or immunotherapy. Radiother Oncol. 2025:110910.

5. Breen WG, Paulino AC, Hartsell WF, Mangona VS, Perkins SM, Indelicato DJ, et al. Factors Associated With Acute Toxicity in Pediatric Patients Treated With Proton Radiation Therapy: A Report From the Pediatric Proton Consortium Registry. Pract Radiat Oncol. 2022;12(2):155–62.

6. Ramseier JY, Ferreira MN, Leventhal JS. Dermatologic toxicities associated with radiation therapy in women with breast cancer. Int J Womens Dermatol. 2020;6(5):349–56.

7. Zhang Q, Gerweck LE, Cascio E, Yang Q, Huang P, Niemierko A, et al. Proton FLASH effects on mouse skin at different oxygen tensions. Phys Med Biol. 2023;68(5).

8. Favaudon V, Caplier L, Monceau V, Pouzoulet F, Sayarath M, Fouillade C, et al. Ultrahigh dose-rate FLASH irradiation increases the differential response between normal and tumor tissue in mice. Sci Transl Med. 2014;6(245):245ra93.

9. Liljedahl E, Konradsson E, Gustafsson E, Jonsson KF, Olofsson JK, Ceberg C, et al. Long-term anti-tumor effects following both conventional radiotherapy and FLASH in fully immunocompetent animals with glioblastoma. Sci Rep. 2022;12(1):12285.

10. Montay-Gruel P, Acharya MM, Goncalves Jorge P, Petit B, Petridis IG, Fuchs P, et al. Hypofractionated FLASH-RT as an Effective Treatment against Glioblastoma that Reduces Neurocognitive Side Effects in Mice. Clin Cancer Res. 2021;27(3):775–84.

11. Velalopoulou A, Karagounis IV, Cramer GM, Kim MM, Skoufos G, Goia D, et al. FLASH Proton Radiotherapy Spares Normal Epithelial and Mesenchymal Tissues While Preserving Sarcoma Response. Cancer Res. 2021;81(18):4808–21.

12. Vozenin MC, De Fornel P, Petersson K, Favaudon V, Jaccard M, Germond JF, et al. The Advantage of FLASH Radiotherapy Confirmed in Mini-pig and Cat-cancer Patients. Clin Cancer Res. 2019;25(1):35–42.

13. Grilj V, Zayas AV, Sesink A, Devanand P, Repáraz D, Paisley R, et al. Average Dose Rate is the Major Temporal Beam Structure Parameter for Preserving Murine Intestines with Pulsed Electron FLASH-RT. Int J Radiat Oncol Biol Phys. 2025.

14. Kim K, Kim MM, Skoufos G, Diffenderfer ES, Motlagh SAO, Kokkorakis M, et al. FLASH Proton Radiation Therapy Mitigates Inflammatory and Fibrotic Pathways and Preserves Cardiac Function in a Preclinical Mouse Model of Radiation-Induced Heart Disease. Int J Radiat Oncol Biol Phys. 2024;119(4):1234–47.

15. Levy K, Natarajan S, Wang J, Chow S, Eggold JT, Loo PE, et al. Abdominal FLASH irradiation reduces radiation-induced gastrointestinal toxicity for the treatment of ovarian cancer in mice. Sci Rep. 2020;10(1):21600.

16. Sorensen BS, Kanouta E, Ankjaergaard C, Kristensen L, Johansen JG, Sitarz MK, et al. Proton FLASH: Impact of Dose Rate and Split Dose on Acute Skin Toxicity in a Murine Model. Int J Radiat Oncol Biol Phys. 2024;120(1):265–75.

17. Almeida A, Godfroid C, Leavitt RJ, Montay-Gruel P, Petit B, Romero J, et al. Antitumor Effect by Either FLASH or Conventional Dose Rate Irradiation Involves Equivalent Immune Responses. Int J Radiat Oncol Biol Phys. 2024;118(4):1110–22.

18. Sorensen BS, Sitarz MK, Ankjaergaard C, Johansen JG, Andersen CE, Kanouta E, et al. Pencil beam scanning proton FLASH maintains tumor control while normal tissue damage is reduced in a mouse model. Radiother Oncol. 2022;175:178–84.

19. Ni H, Reitman ZJ, Zou W, Akhtar MN, Paul R, Huang M, et al. FLASH radiation reprograms lipid metabolism and macrophage immunity and sensitizes medulloblastoma to CAR-T cell therapy. Nat Cancer. 2025;6(3):460–73.

20. Montay-Gruel P, Petersson K, Jaccard M, Boivin G, Germond JF, Petit B, et al. Irradiation in a flash: Unique sparing of memory in mice after whole brain irradiation with dose rates above 100Gy/s. Radiother Oncol. 2017;124(3):365–9.

21. Soto LA, Casey KM, Wang J, Blaney A, Manjappa R, Breitkreutz D, et al. FLASH Irradiation Results in Reduced Severe Skin Toxicity Compared to Conventional-Dose-Rate Irradiation. Radiat Res. 2020;194(6):618–24.

22. Montay-Gruel P, Markarian M, Allen BD, Baddour JD, Giedzinski E, Jorge PG, et al. Ultra-High-Dose-Rate FLASH Irradiation Limits Reactive Gliosis in the Brain. Radiat Res. 2020;194(6):636–45.

23. Valdes Zayas A, Kumari N, Liu K, Neill D, Delahoussaye A, Goncalves Jorge P, et al. Independent Reproduction of the FLASH Effect on the Gastrointestinal Tract: A Multi-Institutional Comparative Study. Cancers (Basel). 2023;15(7).

24. Bohlen TT, Germond JF, Bourhis J, Vozenin MC, Ozsahin EM, Bochud F, et al. Normal Tissue Sparing by FLASH as a Function of Single-Fraction Dose: A Quantitative Analysis. Int J Radiat Oncol Biol Phys. 2022;114(5):1032–44.

25. Kristensen L, Rohrer S, Hoffmann L, Praestegaard LH, Ankjaergaard C, Andersen CE, et al. Electron vs proton FLASH radiation on murine skin toxicity. Radiother Oncol. 2025;206:110796.

26. Ruan JL, Lee C, Wouters S, Tullis IDC, Verslegers M, Mysara M, et al. Irradiation at Ultra-High (FLASH) Dose Rates Reduces Acute Normal Tissue Toxicity in the Mouse Gastrointestinal System. Int J Radiat Oncol Biol Phys. 2021;111(5):1250–61.

27. Allen BD, Alaghband Y, Kramar EA, Ru N, Petit B, Grilj V, et al. Elucidating the neurological mechanism of the FLASH effect in juvenile mice exposed to hypofractionated radiotherapy. Neuro Oncol. 2023;25(5):927–39.

28. Alaghband Y, Allen BD, Kramar EA, Zhang R, Drayson OGG, Ru N, et al. Uncovering the Protective Neurologic Mechanisms of Hypofractionated FLASH Radiotherapy. Cancer Res Commun. 2023;3(4):725–37.

29. Limoli CL, Kramar EA, Almeida A, Petit B, Grilj V, Baulch JE, et al. The sparing effect of FLASH-RT on synaptic plasticity is maintained in mice with standard fractionation. Radiother Oncol. 2023;186:109767.

30. Moeckli R, Goncalves Jorge P, Grilj V, Oesterle R, Cherbuin N, Bourhis J, et al. Commissioning of an ultra-high dose rate pulsed electron beam medical LINAC for FLASH RT preclinical animal experiments and future clinical human protocols. Med Phys. 2021;48(6):3134–42.

31. Marinelli M, Felici G, Galante F, Gasparini A, Giuliano L, Heinrich S, et al. Design, realization, and characterization of a novel diamond detector prototype for FLASH radiotherapy dosimetry. Med Phys. 2022;49(3):1902–10.

32. Jaccard M, Petersson K, Buchillier T, Germond JF, Duran MT, Vozenin MC, et al. High dose-per-pulse electron beam dosimetry: Usability and dose-rate independence of EBT3 Gafchromic films. Med Phys. 2017;44(2):725–35.

33. Verginadis, II, Velalopoulou A, Kim MM, Kim K, Paraskevaidis I, Bell B, et al. FLASH proton reirradiation, with or without hypofractionation, reduces chronic toxicity in the normal murine intestine, skin, and bone. Radiother Oncol. 2025;205:110744.

34. Bohlen TT, Germond JF, Bourhis J, Bailat C, Bochud F, Moeckli R. The minimal FLASH sparing effect needed to compensate the increase of radiobiological damage due to hypofractionation for late-reacting tissues. Med Phys. 2022;49(12):7672–82.

35. Widmark A, Gunnlaugsson A, Beckman L, Thellenberg-Karlsson C, Hoyer M, Lagerlund M, et al. Ultra-hypofractionated versus conventionally fractionated radiotherapy for prostate cancer: 5-year outcomes of the HYPO-RT-PC randomised, non-inferiority, phase 3 trial. Lancet 2019; 394(10196):385–395.

36. Herrera FG, Valerio M, Berthold D, Tawadros T, Meuwly JY, Vallet V, et al. 50-Gy Stereotactic Body Radiation Therapy to the Dominant Intraprostatic Nodule: Results From a Phase 1a/b Trial. Int J Radiat Oncol Biol Phys 2019, 103(2):320:334.

