## Supplementary Materials for "Decrease in dose per fraction impairs the FLASH sparing effect in murine intestine model"

Table S1. Beam parameters used for single-fraction FLASH-RT irradiations.

| Prescribed fraction dose (Gy) | 14 – 20 |
| --- | --- |
| Dose per pulse (Gy) | 2 |
| Pulse width (µs) | 1.9 |
| Dose rate in the pulse (Gy/s) | 1.1E6 |
| Number of pulses | 7 – 10 |
| Pulse repetition frequency (Hz) | 90 |
| Treatment time (ms) | 67 – 100 |
| Average dose rate (Gy/s) | 202 - 209 |

*Table S2. Beam parameters used for two-fraction FLASH-RT irradiations.*

| Prescribed fraction dose (Gy) | 8 – 13 |
| --- | --- |
| Dose per pulse (Gy) | 2 – 2.25 |
| Pulse width (µs) | 1.79 – 1.98 |
| Dose rate in the pulse (Gy/s) | 1.1E6 |
| Number of pulses | 4 – 6 |
| Pulse repetition frequency (Hz) | 90 |
| Treatment time (ms) | 33 – 55 |
| Average dose rate (Gy/s) | 240 - 270 |

*Table S3. Beam parameters used for ten-fraction FLASH-RT irradiations.*

| Prescribed dose (Gy) | 5 – 7 |
| --- | --- |
| Dose per pulse (Gy) | 2.5 – 3.5 |
| Pulse width (µs) | 2.24 – 3.20 |
| Dose rate in the pulse (Gy/s) | 1.1E6 |
| Number of pulses | 2 |
| Pulse repetition frequency (Hz) | 90 |
| Treatment time (ms) | 11 |
| Average dose rate (Gy/s) | 455 - 636 |

| Prescribed fraction dose (Gy) | 5 – 17 |
| --- | --- |
| Dose per pulse (Gy) | 0.02 |
| Pulse width (µs) | 1.2 |
| Dose rate in the pulse (Gy/s) | 16E3 |
| Number of MU | 144 – 488 |
| Pulse repetition frequency (Hz) | 30 |
| Treatment time (s) | 8 – 35 |
| Average dose rate (Gy/s) | 0.5 |

*Table S4. Beam parameters used for CONV irradiations.*

|  | flashDiamond | |
| --- | --- | --- |
| Prescribed fraction dose (Gy) | CONV | FLASH |
| 14 | 13.98 $\pm$ 0.09 | 13.89 $\pm$ 0.14 |
| 15 | 14.99 $\pm$ 0.09 |  |
| 16 | 15.93 $\pm$ 0.07 | 15.93 $\pm$ 0.18 |
| 17 | 17.06 $\pm$ 0.07 | 16.98 $\pm$ 0.13 |
| 18 |  | 17.75 $\pm$ 0.27 |
| 20 |  | 19.95 $\pm$ 0.11 |

*Table S5. Results of dose verification measurements for single-fraction experiments obtained with flashDiamond in water equivalent RW3 slab phantom.*

*Table S6. Results of dose verification measurements for two-fraction experiments obtained with flashDiamond in water equivalent RW3 slab phantom*

|  | flashDiamond | |
| --- | --- | --- |
| Prescribed fraction dose (Gy) | CONV | FLASH |
| 8 | 8.00 $\pm$ 0.02 | 8.08 $\pm$ 0.05 |
| 9 | 8.91 $\pm$ 0.02 | 8.92 $\pm$ 0.05 |
| 10 | 9.93 $\pm$ 0.02 | 10.16 $\pm$ 0.08 |
| 11 | 10.94 $\pm$ 0.01 | 10.98 $\pm$ 0.03 |
| 12 | 11.93 $\pm$ 0.05 | 12.1 $\pm$ 0.1 |
| 13 | 12.92 $\pm$ 0.05 | 12.94 $\pm$ 0.02 |

*Table S7. Results of dose verification measurements for ten-fraction experiments obtained with flashDiamond in water equivalent RW3 slab phantom*

|  | flashDiamond | |
| --- | --- | --- |
| Prescribed fraction dose (Gy) | CONV | FLASH |
| 5 | 4.98 $\pm$ 0.03 | 5.00 $\pm$ 0.05 |
| 5.5 | 5.47 $\pm$ 0.03 | 5.50 $\pm$ 0.05 |
| 6 | 5.99 $\pm$ 0.04 | 6.01 $\pm$ 0.05 |
| 6.5 | 6.49 $\pm$ 0.02 | 6.56 $\pm$ 0.04 |
| 7 | 6.97 $\pm$ 0.03 | 7.01 $\pm$ 0.04 |

*Table S8. Median survival times for single-fraction treatments. Nr stand for not reached.*

|  | Median survival time (days) | |
| --- | --- | --- |
| Fraction dose (Gy) | CONV | FLASH |
| 14 | nr | nr |
| 15 | nr |  |
| 16 | 8 | nr |
| 17 | 6 | 11.5 |
| 18 |  | 8 |
| 20 |  | 7 |

*Table S9. Median survival times for two-fraction treatments. Nr stand for not reached.*

|  | Median survival time (days) | |
| --- | --- | --- |
| Fraction dose (Gy) | CONV | FLASH |
| 8 | nr | nr |
| 9 | nr | nr |
| 10 | 8 | nr |
| 11 | 7 | 8 |
| 12 | 7 | 6 |
| 13 | 7 | 6 |

*Table S10. Median survival times for ten-fraction treatments. Nr stand for not reached.*

|  | Median survival time (days) | |
| --- | --- | --- |
| Fraction dose (Gy) | CONV | FLASH |
| 5 | nr | nr |
| 5.5 | nr | nr |
| 6 | 15 | 12 |
| 6.5 | 9 | 9 |
| 7 | 8 | 8 |

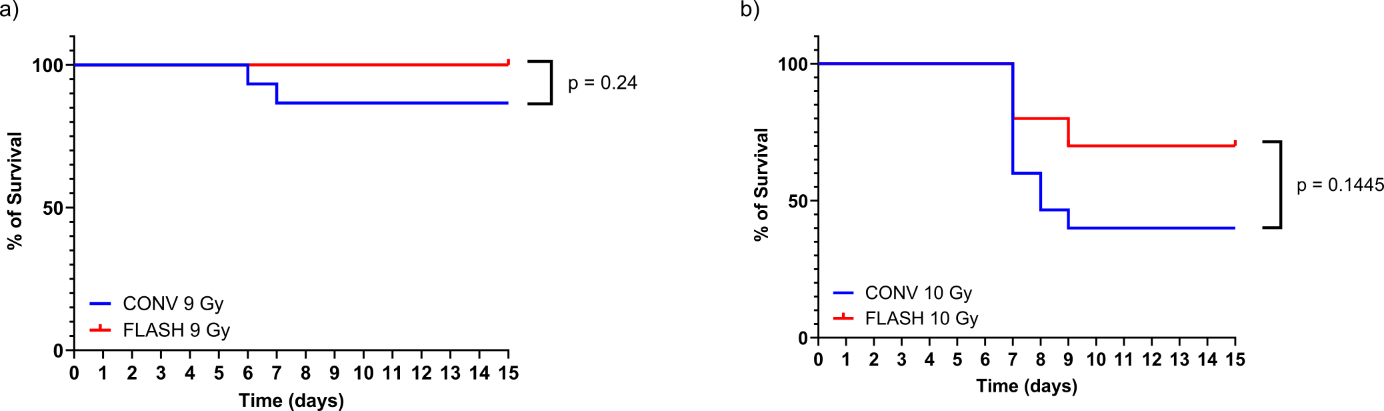

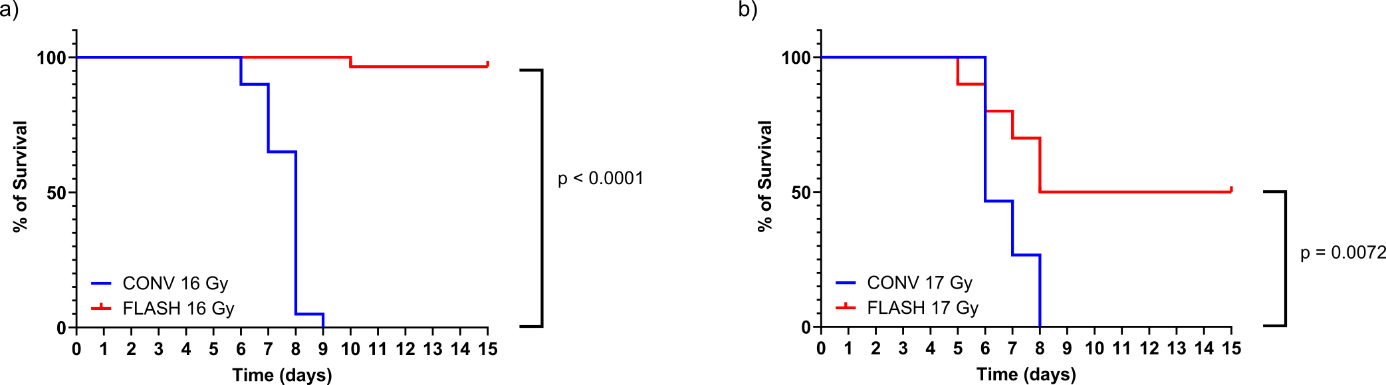

**Supplementary Figure 1.** FLASH confers significant survival advantage over CONV at 16 and 17 Gy single fractions. a) Kaplan–Meier survival curves showing 15-day survival following single-fraction whole-abdomen irradiation with 16 Gy CONV (n = 20) or 16 Gy FLASH (n = 29). Survival at day 15 was 0% in the CONV group and 95.6% in the FLASH group (*p* < 0.0001). b) Kaplan–Meier survival curves showing 15-day survival following single-fraction whole-abdomen irradiation with 17 Gy CONV (n = 15) or 17 Gy FLASH (n = 10). Survival at day 15 was 0% in the CONV group and 50% in the FLASH group (*p* = 0.0072).

**Supplementary Figure 2.** Non-significant survival advantage of FLASH over CONV in two-fraction regimens at 9 and 10 Gy per fraction. a) Kaplan–Meier survival curves showing 15-day survival following whole-abdomen irradiation with two fractions of 9 Gy delivered by CONV (n = 15) or FLASH (n = 10). Survival rates were 86.7% in the CONV group and 100% in the FLASH group (*p* = 0.24). b) Kaplan–Meier survival curves showing 15-day survival following whole-abdomen irradiation with two fractions of 10 Gy delivered by CONV (n = 15) or FLASH (n = 10). The survival rates observed were 40% in the CONV group and 70% in the FLASH group (*p* = 0.1445).

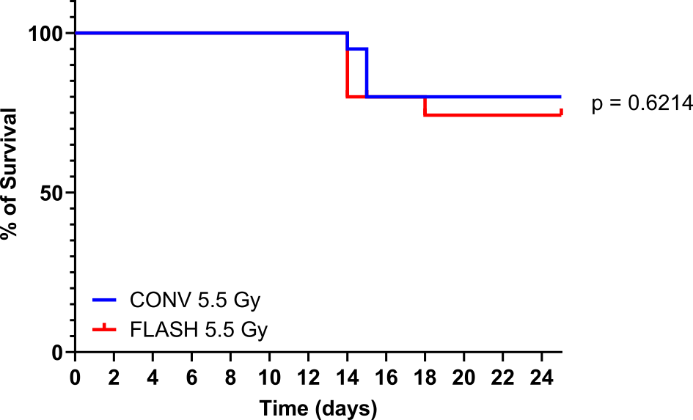

**Supplementary Figure 3.** No survival benefit of FLASH over CONV in a ten-fraction regimen at 5.5 Gy per fraction. Kaplan–Meier survival curves showing 25-day survival following whole-abdomen irradiation with ten fractions of 5.5 Gy delivered by CONV (n = 20) or FLASH (n = 20). The survival rates observed for were 80% in the CONV group and 75% in the FLASH group (*p* = 0.6214).
